# Individual gait is not the optimum of standard cost functions: a test with two predictive simulation frameworks

**DOI:** 10.64898/2026.11.24.768861

**Authors:** Muhammad Mughees Ahmed, David Muñoz, Giacomo Severini

**Affiliations:** School of Electrical and Electronic Engineering, University College Dublin, Belfield, Dublin 4, D04 V1W8, Ireland; Brussels Human Robotics Research Center (BruBotics), Vrije Universiteit Brussel, Pleinlaan 2, 1050 Brussels, Belgium

**Keywords:** Predictive simulation, Gait, Inverse optimal control, Cost function, Optimality, Musculoskeletal modelling, Reflex control, Direct collocation

## Abstract

**Background:** Predictive simulations of gait are increasingly proposed as a way to predict how individuals, including people with impaired gait, will walk. Most approaches generate gait by minimising a cost function, a weighted sum of terms such as metabolic energy and muscle activation. The weights of the cost can be personalised using inverse optimal control (IOC), but the basic assumption that an individual’s gait is the optimum of a cost is rarely questioned or tested.

**Methods:** We tested this assumption on the openly available data of healthy adults with subject-scaled models, using a shooting framework with a reflex controller (implemented in SCONE) and a direct collocation framework (implemented in PredSim). For each individual, we compared a simulation that tracked the measured joint angles, electromyography and stride time with the gaits that minimised the framework’s standard cost at 35 weightings. We repeated the comparison for averages of 10 to 30 subjects. Paired comparisons across the 20 subjects used the Wilcoxon signed-rank test, and averages were compared with the mean of their members by generalised least squares.

**Results:** In both frameworks, the standard-cost optimum tracked the data worse than the tracked gait in every subject, and the tracked gait always had a higher cost. Reweighting lowered the tracking error of the optimum, but the best weighting still tracked the data worse than the tracked gait in both frameworks. When the data and bodies of subjects were averaged, the standard-cost optimum of the average tracked the averaged data 30% more closely than individual optima tracked individual data.

**Conclusions:** Changing the weights of cost function elements only selects a different optimum of the same cost terms, whereas individual gait, as represented by the tracked gait, was not an optimum of the standard cost. The closer match between optima and averaged data suggests that comparisons with group data overstate how well standard costs describe individuals. Predicting individual gait therefore requires formulations that allow the simulated gait to deviate from the optimum of a cost.

## Background

A common approach in predictive simulation is to generate gait by optimising the control of a musculoskeletal model, so that the movement results from the minimisation of a cost function, usually a weighted sum of metabolic energy, muscle activation and a term for smoothness or stability [1–5].

Among the optimality-based approaches, shooting methods use the optimisation to select the parameters of a neural controller that drives a forward simulation [6–8]. Direct collocation methods minimise the cost directly over the states and controls of the gait [3, 9]. In both approaches, the cost terms and their weights are usually defined a priori and kept fixed through the different scenarios that are predicted. A cost is usually judged by how closely the predicted gait matches measured gait, and a close match is often read as evidence that people walk by minimising that cost. These comparisons have mostly used generic models and group-average data, which represent an average person rather than a specific individual.

Predictive simulations are increasingly proposed as a tool to generate predictions for individuals with impaired gait. Falisse et al. [10] predicted the crouch gait of one child with cerebral palsy using a model built from medical images. Van Den Bosch et al. [11] built personalised models of eight children with cerebral palsy and found that the effect of each impairment on gait differed between children. In people with plantarflexor weakness, the ankle-foot orthosis stiffness that minimised walking energy cost depended on the severity of the weakness [12], and Kiss et al. [13] proposed using such simulations to prescribe orthoses for individual patients. All these applications assume that minimising a cost function predicts how a specific, impaired person walks.

In predictive simulations of gait, the principle of optimality itself is rarely questioned. Where the field has addressed individual differences, it has done so by personalising the weights of the cost function. To this aim, inverse optimal control (IOC) is used to search for the weighting of the cost terms whose optimally generated gait best tracks the measured data [14]. Identified weights differ between subjects [15] and with walking speed [16], but IOC still assumes that the measured gait is an optimum of the cost for some weighting of the chosen terms. What remains untested is whether optimisation can describe an individual’s gait at all, although experimental studies give reasons to doubt it. Gait patterns are unique to each person [17]. People re-optimise their energy cost when walking conditions change [18], but they do not always converge right away on the cheapest gait [19, 20]. When a robot perturbed their gait, people adapted to perturbations that challenged stability even at a higher metabolic cost and ignored those that did not [21, 22]. Muscle coordination has also been shown to be habitual rather than optimal [23].

Optimality can be tested, but only up to the limit in precision of the modelling framework. If an individual walks by minimising a cost, a simulation that tracks the person’s data should lie at or near the minimum of that cost, and the gait that minimises the cost should track the person’s data about as closely as the tracking simulation does [5, 14, 24]. In this study, we tested whether the gait of an individual is consistent with the optimisation of a standard cost function. We used two frameworks: a shooting framework with a reflex controller (SCONE) and a direct collocation framework (PredSim). In 20 healthy adults with subject-scaled models, we compared the tracked gait of each subject with the gait that minimised the framework’s standard cost. We did this both at the default weights and at the best of 35 weightings of two cost terms. All gaits were scored with the same tracking error, which combined joint angles, electromyography (EMG) and stride time. Because tracking and prediction used the same model, constraints and solver, the differences between them can be mostly attributed to the formulation of the objective as a cost. Finally, we tested whether the standard cost describes averages of people better than individuals. We hypothesised that (1) the tracked gait of an average, built from averaged data and an averaged body, has a lower standard cost than the tracked gaits of its members, and (2) the standard-cost optimum of an average tracks its data better than the members’ optima track theirs.

## Methods

### Dataset

We used the open gait dataset of Van Criekinge et al. [25], which contains motion capture, ground reaction forces and surface EMG of 138 healthy adults walking overground at their self-selected speed. All analyses used EMG of five muscle groups on both legs: hamstrings (semitendinosus and biceps femoris), rectus femoris, vastus lateralis, gastrocnemius and tibialis anterior. For each subject, we scaled the musculoskeletal model (see Modelling) to the subject’s marker data in OpenSim, then computed hip, knee, ankle and pelvis angles for every stride with OpenSim’s inverse kinematics from the Plug-in-Gait marker trajectories. For the analysis of individuals in both frameworks, we selected 20 subjects with clean inverse kinematics data and EMG. Together they covered the dataset’s range of age (about 25 to 87 years), body mass (52 to 93 kg), leg length (0.79 to 1.00 m) and walking speed (0.80 to 1.47 m/s). For the analysis of averages, we used a larger group of 70 subjects: all subjects with EMG of the five muscle groups and usable inverse kinematics (body mass 72 ± 13 kg, leg length 0.88 ± 0.06 m; mean ± SD). This group includes the 20 subjects above. In our analyses we used, for each subject, the hip, knee and ankle flexion angles of both legs, pelvis tilt, the EMG envelopes and the stride time. Each leg was cut on its own strides and resampled to 360 points, and the subject’s data are the mean and standard deviation (SD) across strides. EMG was normalised by dividing its mean over the stride.

### Modelling and evaluation

Both frameworks used the same planar musculoskeletal model (derived from the gait10dof18musc Opensim Model [26]) with 9 degrees of freedom (pelvis tilt and translations, and hip, knee and ankle of each leg, with the trunk locked to the pelvis, the tenth degree of freedom present in the original model, the lumbar joint, is welded in this implementation) and 18 Hill-type muscles (hamstrings, biceps femoris short head, gluteus maximus, iliopsoas, rectus femoris, vasti, gastrocnemius, soleus and tibialis anterior on each side). The model was scaled to each subject: segment dimensions, masses and inertias, joint positions, muscle paths, maximal isometric forces, optimal fibre lengths and tendon slack lengths. The two frameworks used the same scaled muscle parameters. Foot–ground contact was modelled with two spheres per foot (heel and toe), using each framework’s own contact model: Hyfydy’s contact model in SCONE, with sphere positions scaled to the subject’s foot, and smooth sphere–half-space contacts in PredSim [3], with the same sphere positions in all subjects.

We scored every simulated gait with one tracking error, computed in the same way in both frameworks, so that their results could be compared. The two frameworks tracked the data with slightly different objectives: SCONE minimised this tracking error directly, whereas PredSim tracked the mean curves without the band. For each joint angle and EMG channel, the error is the mean squared distance of the simulated curve outside the band of the subject’s mean ± 1 SD (with a minimum band width of 4° for angles and 0.3 for normalised EMG). The total tracking error is 0.1 × the mean tracking error over the four angles squared (deg²) + 10 × the mean tracking error over the five EMG channels + 100 × (*T*/*T*_meas_ − 1)², where *T* is the simulated and *T*_meas_ the measured stride time, averaged over both legs. Lower values of this tracking error mean a closer match. For example, if all four joint angles lie 5° outside the band throughout the stride, the angle part is 0.1 × 25 = 2.5. If one EMG channel lies 1.0 outside its band throughout the stride (that is, a deviation equal to its mean activation) and the other four stay inside, the EMG part is 10 × 1/5 = 2. A stride that is 10% too long or too short adds 100 × 0.1² = 1.

Each framework has its own standard cost function, with terms and default weights taken from the literature and from the framework’s defaults (Table 1). Here we refer to the gait that minimises the standard cost with the default weights as the standard-cost optimum.

**Table 1.** Standard cost of the two frameworks.

| Term | SCONE | PredSim |
|---|---|---|
| Metabolic energy | Cost of transport [27]; weight 1 | Energy rate squared [28];<br>weight 500 |
| Muscle effort (grid rows) | Squared activation;<br>weight 10 | Squared activation;<br>weight 2000 |
| Movement regularisation | Head acceleration;<br>weight 0.25 | Joint accelerations squared;<br>weight 50000 |
| Joint limits | Ankle range and knee limit torque;<br>weights 0.067 and 0.0067 | Passive joint torques squared;<br>weight 1000 |
| Other regularising terms | Vertical ground reaction force above 1.5 body weight;<br>weight 100<br>Falling and speed outside measured ± 0.05 m/s;<br>weight 1000 | Speed fixed to the measured speed (constraint); slack terms;<br>weight 0.001 |

All terms are normalised by distance travelled. The two terms varied in the weight grid are marked.

For each subject and framework, we compared three gaits:

1) the tracked gait, which is the gait obtained by having the framework track the data of an individual as closely as possible
2) the standard-cost optimum, which is the gait that minimises the standard cost function with its default weights
3) the best weighting gait, which is the gait obtained with the weights of the standard cost function optimised (across 35 different combinations) to track the data as closely as possible

Paired comparisons used the Wilcoxon signed-rank test (n = 20), as the differences were not normally distributed.

For the analysis of averages, each group of subjects was represented by an average subject, with its own data and its own body. The data of an average were built from each member’s data. We took the mean of the members’ mean curves for each joint angle and EMG channel, giving each subject the same weight, then renormalised the averaged EMG to a mean of 1 over the stride. The band around the average used for the tracking score was the members’ mean within-subject SD. The speed, the stride time and the timing of the right heel strike were the means of the members’ values. The body of an average was built by averaging, value by value, every numerical parameter of the members’ scaled models: segment masses, inertias and dimensions, joint positions, contact sphere positions, muscle paths, maximal isometric forces, optimal fibre lengths and tendon slack lengths. The scaled models of all subjects have the same structure and differ only in these values, so the average is a valid model of the same type. It was then used exactly like a subject’s model.

### SCONE: tracking and predicting

For all SCONE simulations we used SCONE 2.0.6 [7] with the Hyfydy engine [29] to simulate the model driven by the reflex controller of Geyer and Herr [6]. We optimised the 39 controller parameters with the covariance matrix adaptation evolution strategy (CMA-ES) [30], and each simulation started from a fixed, subject-specific initial state (joint angles and velocities). To track a subject, we minimised the tracking error explained in the “Modelling” section plus SCONE’s usual regularising terms over 6 s of walking, with two random seeds and 1000 generations each. These terms are penalties for falling, for speeds more than 10% away from the measured speed, for high ground reaction forces and for exceeding joint limits, plus metabolic effort at a low weight. The best walking solution was the subject’s tracked gait.

To predict, we minimised the standard cost (Table 1), with a penalty for speeds more than 0.05 m/s from the measured speed. The activation weight of 10 makes the activation term about 2% of the metabolic term, close to the share used by Veerkamp et al. [5]. We repeated the prediction for 35 weightings: the activation weight multiplied by 0, 0.3, 1, 3 or 10, and the head-acceleration weight by 0.1, 0.32, 1, 3.2, 10, 32 or 100. The combination included the default weights. Each optimisation started from the tracked gait’s controller (three seeds, 800 generations, 5 s of walking), and we kept the seed with the lowest cost. All kept gaits were then evaluated over 5 s, and a gait whose pelvis dropped below 0.6 m was counted as a fall and given no tracking error.

The tracked gaits walked on average 7% slower than measured (range −16% to +15%). As a check, we also re-optimised the standard cost with the speed window centred on each tracked gait’s own speed.

### PredSim: tracking and predicting

PredSim [3] solves the prediction problems by direct collocation (CasADi and IPOPT; third-order Radau collocation with 50 mesh intervals) over one periodic stride, with the speed fixed to the measured speed and the stride time kept free. Tracking had two steps. First we tracked the joint angles alone, then we used the found solutions as starting point for the full tracking error (thus including EMGs and stride time) with a band of zero width. During tracking, a constraint placed the model’s left and right heel strikes at the measured instants, so that simulated and measured strides were aligned. The tracking problem also contained the standard cost at 1/1000 of its weights, to prevent the tracked gaits from co-activating the muscles that had no EMG to track. To predict, we minimised PredSim’s standard cost (Table 1) for 35 weightings: the activation weight multiplied by 1, 3, 10, 30 or 100, and the joint-acceleration weight by 0.032, 0.1, 0.32, 1, 3.2, 10 or 32. Also in this case the grid included the default weights. Every prediction used the tracked gait as starting point.

For the analysis of averages, we drew 60 random groups from the 70 subjects described in the Dataset section: 20 groups each of 10, 20 and 30 subjects. Each of the 70 individuals and 60 averages was tracked, and its standard-cost optimum was computed as above.

To test the two hypotheses stated in the Background, we computed, for each average, the relative difference between its value and the mean of its own members’ values. Since averages share members, we estimated the mean difference by generalised least squares, with a covariance that grows with the number of members two averages share, and report 95% confidence intervals (CI).

### Use of generative AI

The authors used Claude (Anthropic) as an AI assistant throughout this work. Under the authors’ direction, it was used to (i) help write and debug the MATLAB, Python and Lua code that set up and ran the SCONE and PredSim simulations and the analysis pipeline; (ii) run the statistical analyses and derive the data used to make the figures; and (iii) edit the manuscript text to improve the English language and readability of the text. The authors designed the study and made all methodological decisions. The authors checked the code, results and references, and revised all text. The authors take full responsibility for the content of this manuscript.

## Results

### Tracking the person versus optimising the standard cost

In both frameworks, the tracked gait of every subject matched the data more closely than the standard-cost optimum did, and had a higher standard cost (Fig. 1).

**Figure 1.**
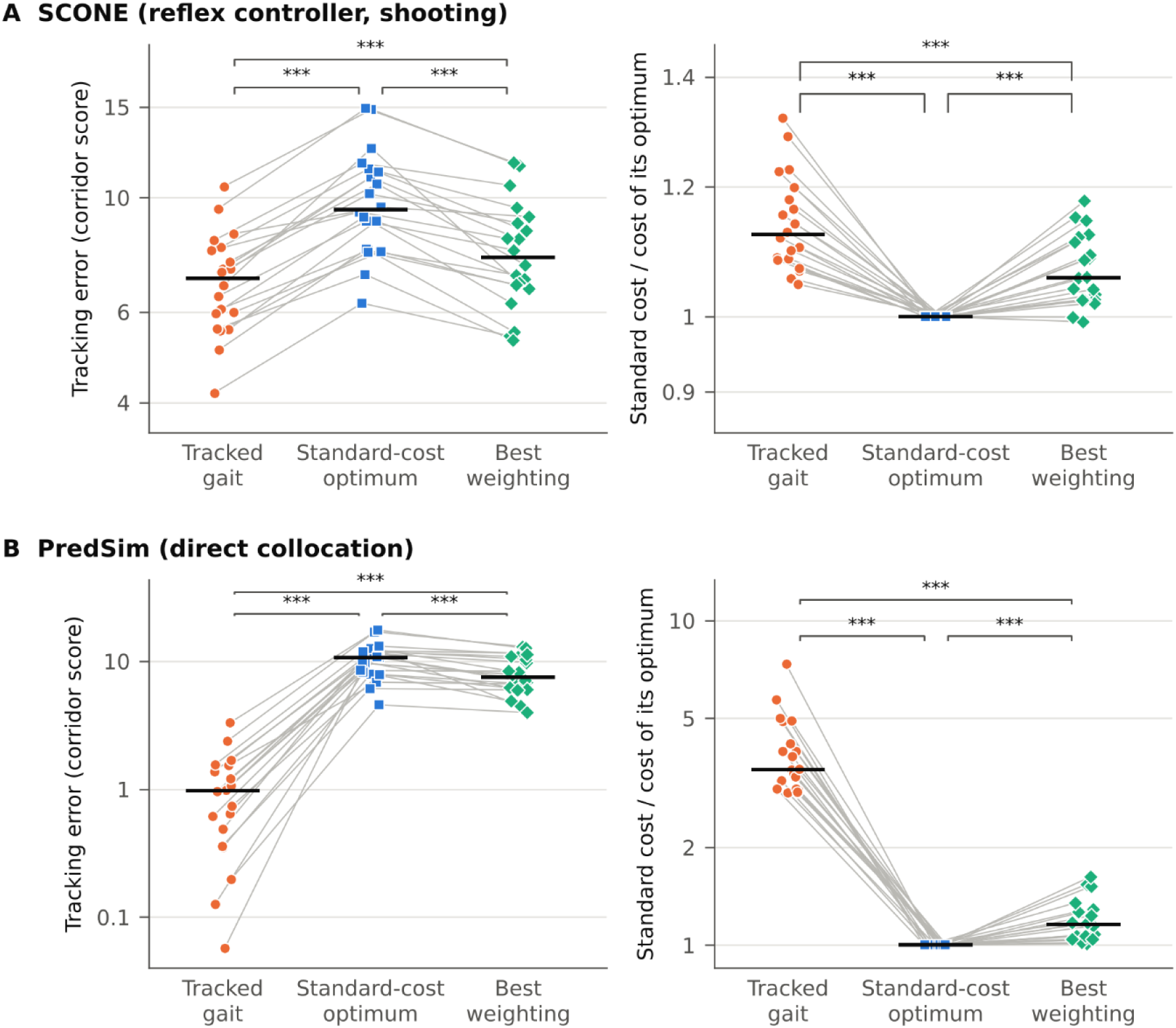
Tracked gait, standard-cost optimum and best weighting in SCONE (A) and PredSim (B). Results for the 20 subjects. Left: tracking error (lower is closer to the subject’s data). Right: standard cost of each gait divided by the cost of the standard-cost optimum. Lines join the same subject; black bars are medians. Vertical axes are logarithmic. Brackets show paired comparisons (Wilcoxon signed-rank test, n = 20; *** p < 0.001).

In SCONE, the median tracking error was 6.96 (range 4.18 to 10.5) for the tracked gaits and 9.48 (6.24 to 14.9) for the standard-cost optima. The standard-cost optimum gait tracked worse in all 20 subjects (Wilcoxon signed-rank test, p < 0.001). The tracked gait cost more in all 20 subjects (p < 0.001). When the speed penalty was centred on the tracked gait’s own speed, the tracked gaits cost 15% more than the optimum (range 7 to 32%).

In PredSim, the median tracking error was 0.97 (range 0.06 to 3.3) for the tracked gaits and 10.7 (4.6 to 17.5) for the standard-cost optima. The standard cost optimum gait tracked worse in all 20 subjects (Wilcoxon signed-rank test, p < 0.001). The standard cost of the tracked gaits was 9005 (median; range 4271 to 21 700), against 2448 (1230 to 4241) for the standard-cost optimum. The tracked gait cost more in all 20 subjects (p < 0.001).

The best weighting from the grid decreased the tracking score of the optimised gait when compared to the standard weighting, but it was still significantly higher than the tracking score of the tracked gait. In SCONE the tracking error of the best weighting was 7.66 (5.29 to 11.7), and was higher than that of the tracked gait in 18 of 20 subjects (median +15%; p < 0.001). The two exceptions tracked slightly better (−1% and −5%). In PredSim it was 7.5 (4.0 to 13.0), higher than that of the tracked gait in all 20 subjects (p < 0.001). The gaits of the best weightings cost little more than the standard-cost optimum and less than the tracked gaits: 6.18 (range 5.47 to 7.20) against 5.79 and 6.65 in SCONE, and 2821 (1896 to 6894) against 2448 and 9005 in PredSim.

### Weight grids

Changing the weights across the grid changed the tracking error, when compared to the default weighting, by −6% to +34% in SCONE, and by −9% to +112% in PredSim (averaged across subjects, Fig. 2).

**Figure 2.**
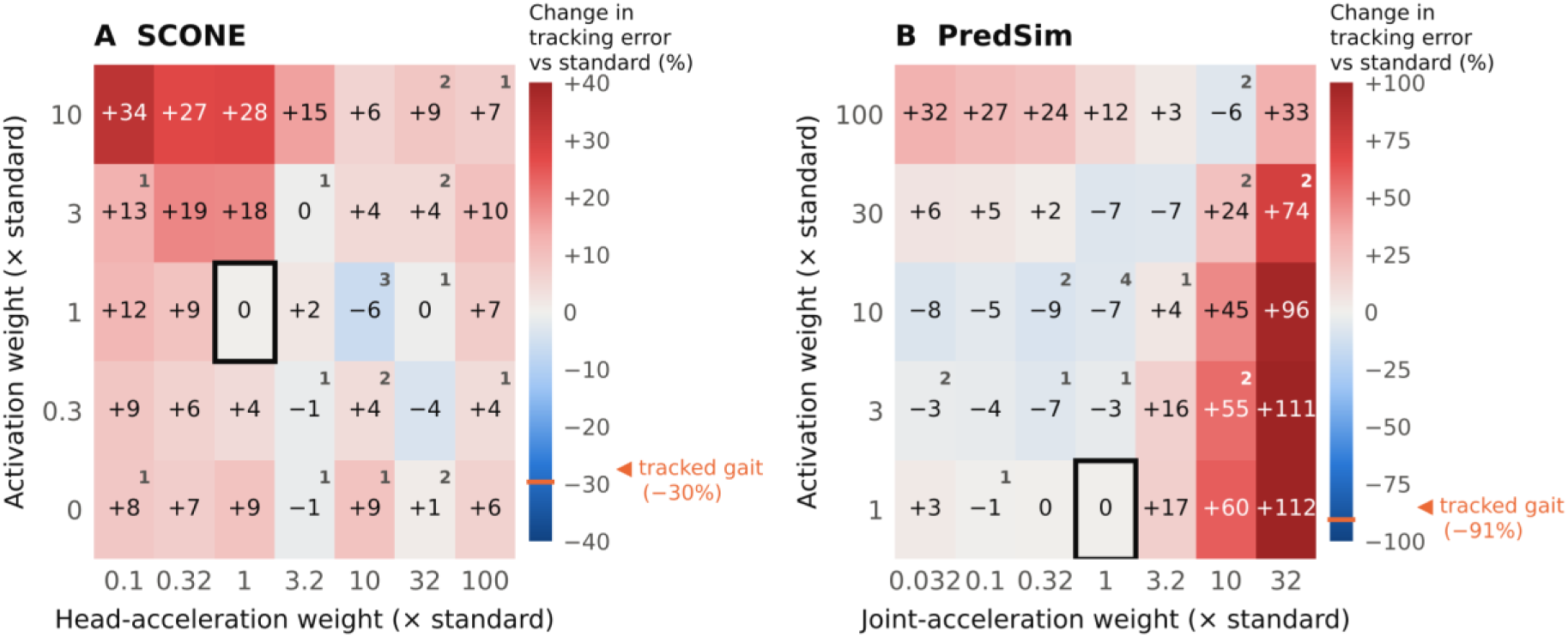
Mean change in tracking error across the 35 weightings in SCONE (A) and PredSim (B). Change relative to the standard weighting (boxed). Blue cells are closer to the data than the standard cost. Means over the 20 subjects. Small numbers give how many of the 20 subjects had their best weighting in that cell. The orange mark on each colour bar is the tracked gait on the same scale.

On the same scale, the tracked gaits had a tracking error that was 30% (SCONE) and 91% (PredSim) below that of the standard-cost optimum. Per subject, the best weighting lowered the tracking error by a median 19% in SCONE and 17% in PredSim. This closed a median 62% (SCONE) and 19% (PredSim) of the distance between the standard-cost optimum and the tracked gait.

The best weighting differed between subjects. It fell in 14 (SCONE) and 11 (PredSim) of the 35 cells, and it was never the standard weighting.

### Joint angles and muscle activity

Tables 2 and 3 compare the three gaits (tracked, standard cost optimum and best optimum) with the measured joint angles, muscle activity and spatiotemporal parameters, and give the elements of the standard cost. Figs. 3 and 4 show a representative subject for each framework, selected as a subject with values of error close to the group average.

**Figure 3.**
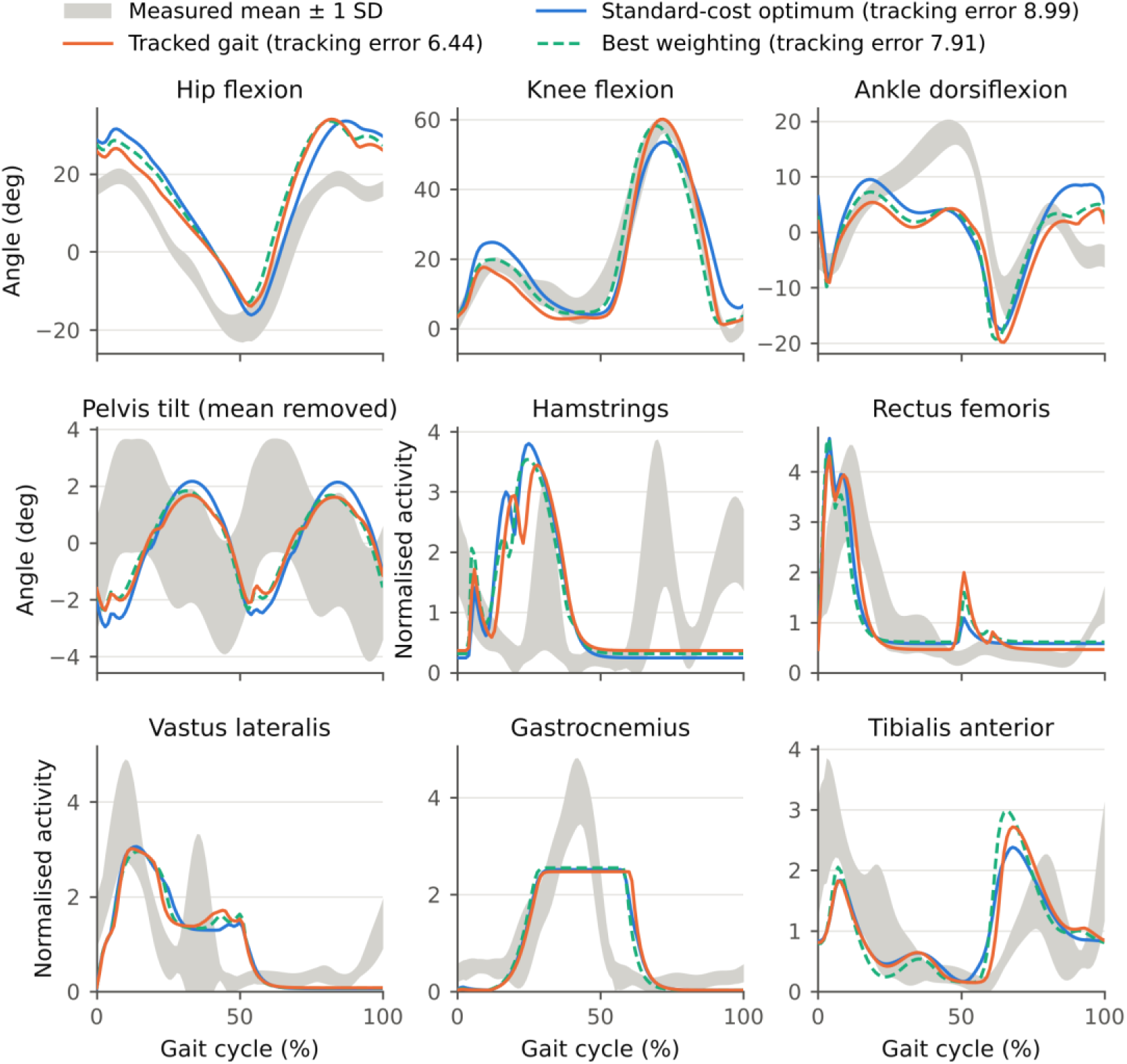
Joint angles and muscle activity of a representative subject in SCONE. Grey bands show the measured mean ± 1 SD; lines show the tracked gait, the standard-cost optimum and the best weighting, with their tracking errors in the legend. Pelvis tilt is shown with its stride mean removed. Muscle activity is the simulated activation and the EMG envelope, each divided by its mean over the stride.

**Figure 4.**
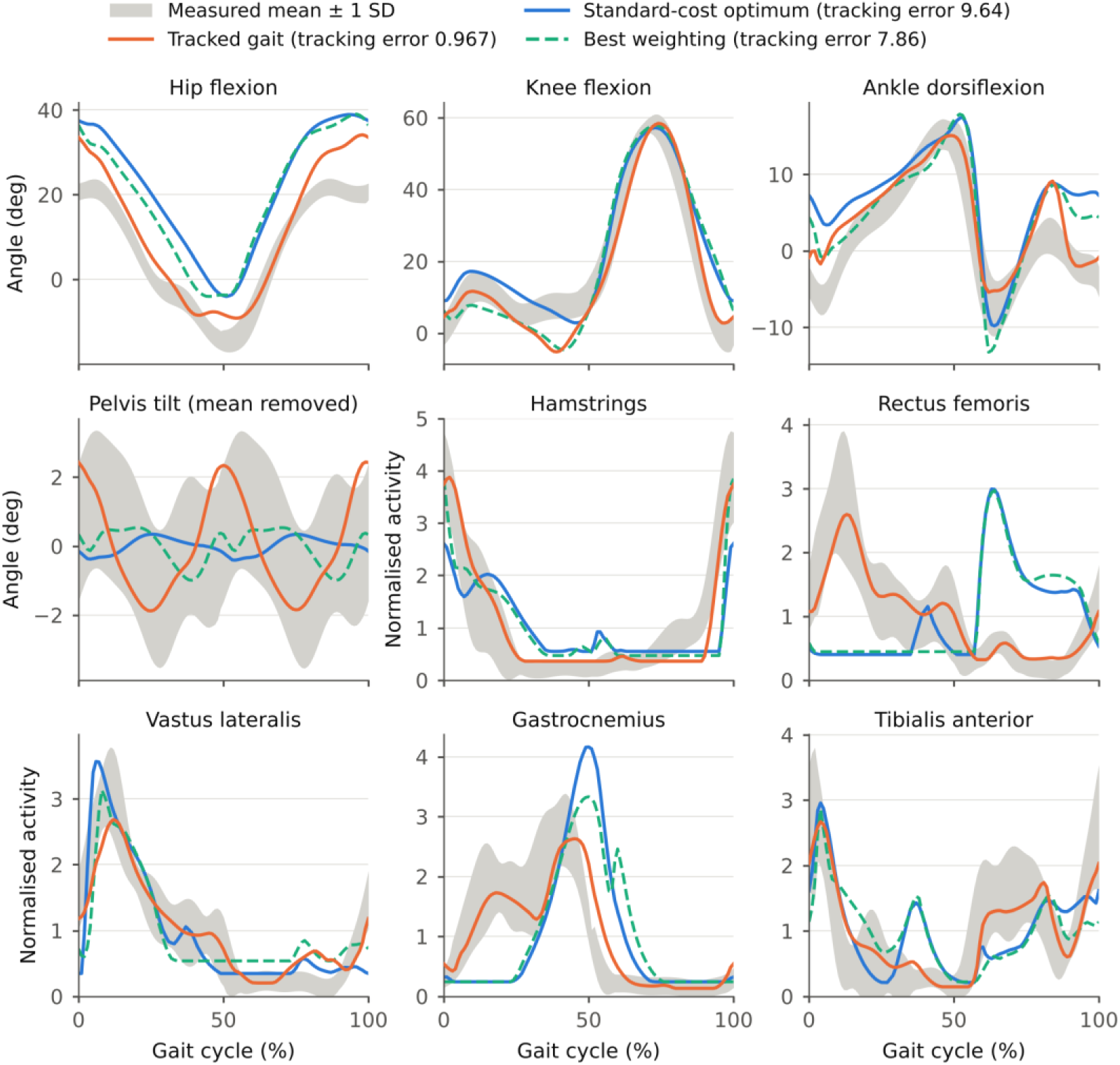
Joint angles and muscle activity of a representative subject in PredSim. Subject 17, one of the subjects whose tracking errors for the three gaits were closest to the group medians. Same layout as Fig. 3.

**Table 2.** Agreement with measured data and standard-cost elements of the three gaits in SCONE.

|  | Tracked gait | Standard-cost optimum | Best weighting |
|---|---|---|---|
| **Joint angles (hip, knee, ankle)** |  |  |  |
| Correlation, r | 0.91 [0.89, 0.92] | 0.88 [0.85, 0.91] | 0.89 [0.87, 0.92] |
| NRMSE (%) | 17.8 [14.8, 22.3] | 20.0 [17.9, 24.0] | 19.3 [16.3, 23.6] |
| **Pelvis tilt** |  |  |  |
| Correlation, r | 0.62 [−0.03, 0.75] | 0.42 [−0.09, 0.58] | 0.53 [−0.08, 0.68] |
| NRMSE (%) | 26.3 [23.4, 52.2] | 32.6 [27.3, 51.3] | 27.8 [23.9, 47.9] |
| **Muscle activity (5 muscles)** |  |  |  |
| Correlation, r | 0.47 [0.39, 0.51] | 0.47 [0.41, 0.51] | 0.43 [0.38, 0.48] |
| NRMSE (%) | 31.9 [30.1, 35.3] | 33.3 [31.0, 37.4] | 32.7 [31.0, 36.0] |
| **Spatiotemporal parameters** |  |  |  |
| Stride time, absolute error (%) | 2.5 [1.3, 4.3] | 5.6 [2.2, 7.9] | 4.7 [2.1, 6.3] |
| Speed, absolute error (%) | 9.7 [7.3, 11.3] | 6.4 [2.7, 8.0] | 8.3 [7.2, 9.4] |
| **Standard cost (weighted elements)** |  |  |  |
| Speed penalty (×1000) | 61.73 [0.00, 87.66] | 0.00 [0.00, 0.00] | 0.00 [0.00, 0.00] |
| Metabolic energy (×1) | 5.65 [5.34, 5.97] | 5.09 [4.94, 5.29] | 5.49 [5.27, 5.62] |
| Muscle activation (×10) | 0.140 [0.114, 0.171] | 0.110 [0.098, 0.122] | 0.136 [0.109, 0.157] |
| Head acceleration (×0.25) | 0.584 [0.519, 0.703] | 0.592 [0.529, 0.677] | 0.487 [0.448, 0.570] |
| Joint limits (×0.067, ×0.0067) | 0.000 [0.000, 0.000] | 0.000 [0.000, 0.000] | 0.000 [0.000, 0.000] |
| Ground reaction force (×100) | 0.177 [0.049, 0.279] | 0.013 [0.005, 0.021] | 0.015 [0.004, 0.128] |
| Total without speed penalty | 6.65 [6.24, 6.97] | 5.79 [5.65, 6.01] | 6.18 [6.00, 6.56] |
| Total | 68.64 [6.92, 94.28] | 5.79 [5.65, 6.01] | 6.18 [6.00, 6.56] |

Tracked gait, standard-cost optimum and best weighting in SCONE (n = 20). Values are medians [interquartile range] across subjects. NRMSE, normalised root-mean-square error.

**Table 3.**
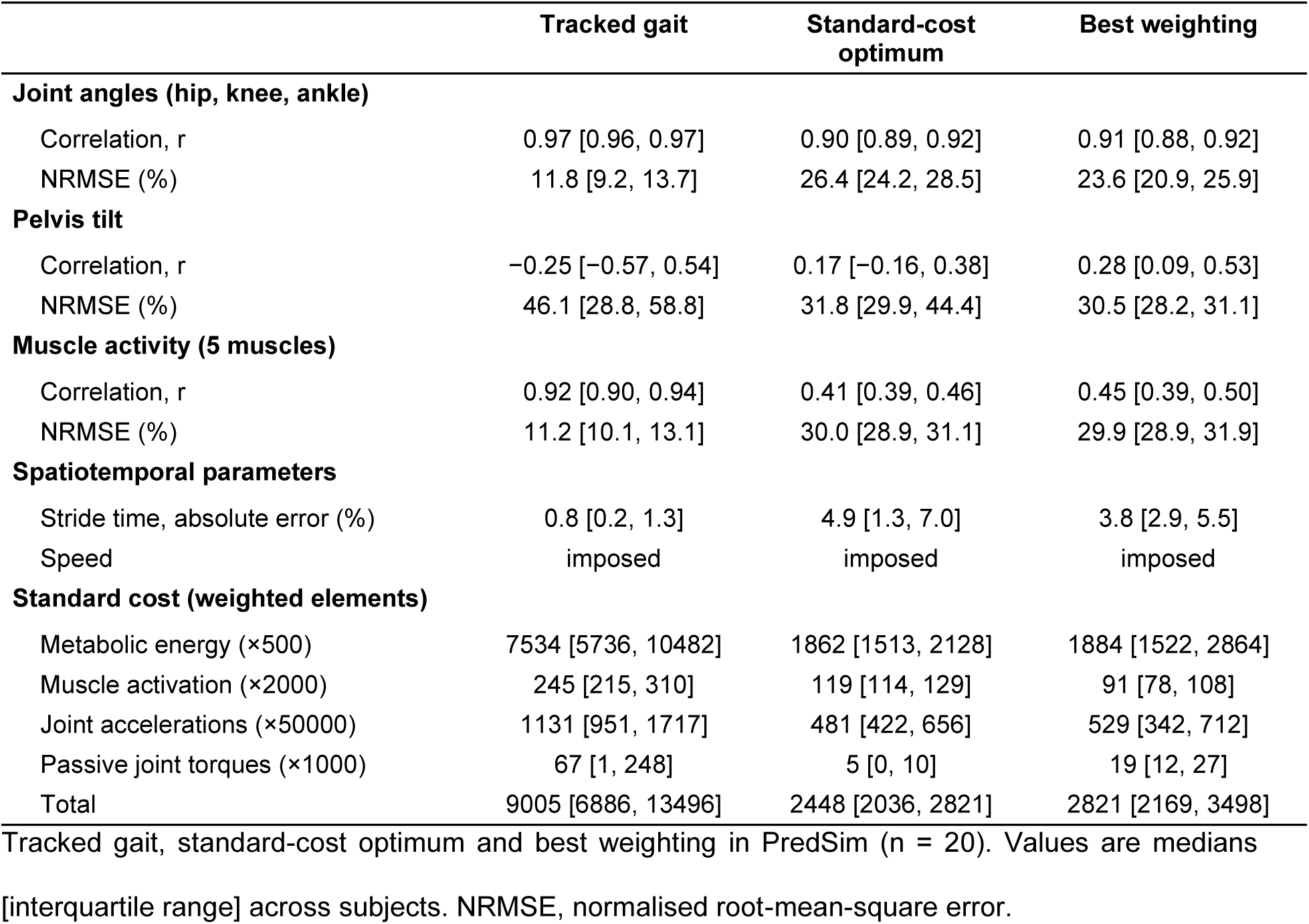
Agreement with measured data and standard-cost elements of the three gaits in PredSim.

|  | **Tracked gait** | **Standard-cost optimum** | **Best weighting** |
|---|---|---|---|
| **Joint angles (hip, knee, ankle)** |  |  |  |
| Correlation, r | 0.97 [0.96, 0.97] | 0.90 [0.89, 0.92] | 0.91 [0.88, 0.92] |
| NRMSE (%) | 11.8 [9.2, 13.7] | 26.4 [24.2, 28.5] | 23.6 [20.9, 25.9] |
| **Pelvis tilt** |  |  |  |
| Correlation, r | −0.25 [−0.57, 0.54] | 0.17 [−0.16, 0.38] | 0.28 [0.09, 0.53] |
| NRMSE (%) | 46.1 [28.8, 58.8] | 31.8 [29.9, 44.4] | 30.5 [28.2, 31.1] |
| **Muscle activity (5 muscles)** |  |  |  |
| Correlation, r | 0.92 [0.90, 0.94] | 0.41 [0.39, 0.46] | 0.45 [0.39, 0.50] |
| NRMSE (%) | 11.2 [10.1, 13.1] | 30.0 [28.9, 31.1] | 29.9 [28.9, 31.9] |
| **Spatiotemporal parameters** |  |  |  |
| Stride time, absolute error (%) | 0.8 [0.2, 1.3] | 4.9 [1.3, 7.0] | 3.8 [2.9, 5.5] |
| Speed | imposed | imposed | imposed |
| **Standard cost (weighted elements)** |  |  |  |
| Metabolic energy (×500) | 7534 [5736, 10482] | 1862 [1513, 2128] | 1884 [1522, 2864] |
| Muscle activation (×2000) | 245 [215, 310] | 119 [114, 129] | 91 [78, 108] |
| Joint accelerations (×50000) | 1131 [951, 1717] | 481 [422, 656] | 529 [342, 712] |
| Passive joint torques (×1000) | 67 [1, 248] | 5 [0, 10] | 19 [12, 27] |
| Total | 9005 [6886, 13496] | 2448 [2036, 2821] | 2821 [2169, 3498] |

Tracked gait, standard-cost optimum and best weighting in PredSim (n = 20). Values are medians [interquartile range] across subjects. NRMSE, normalised root-mean-square error.

In SCONE, the three gaits differed little from each other in joint angles and muscle activity (Table 2, Fig. 3). The tracked gait followed the joint angles slightly more closely than the two optima, but none of the three gaits followed the measured EMG closely. Pelvis tilt was matched less well than the other angles by all three gaits, with the tracked gait the closest of the three. The tracked gaits matched the measured stride time more closely than the optima, but deviated more from the measured speed. The tracked gait had higher metabolic energy, muscle activation and ground reaction force terms than the optima.

In PredSim, the tracked gait followed both the joint angles and the EMG very closely (Table 3, Fig. 4). The standard-cost optimum and the best weighting kept joint-angle patterns similar to the measured ones, but with larger errors, mainly associated with a more flexed hip. The muscle activity of the two optimum gaits correlated poorly with the EMG, and in both the rectus femoris was active in swing instead of around heel strike. Pelvis tilt, which changes by only a few degrees over the stride, was the only angle that the tracked gait matched less closely than the optima. The optima also deviated more from the measured stride time. In 16 of 20 subjects, the tracked gait had higher values than the standard-cost optimum for all four elements of the standard cost (14 of 20 against the best weighting), with the largest difference observed in metabolic energy.

### Averages analysis in PredSim

Averaging the data and bodies across multiple participants did not make the tracked gait cheaper (Fig. 5). The standard cost of the averages’ tracked gaits was 5.0% higher than that of their members (95% CI −2.5 to +12.5%; p = 0.19), and the result did not change with group size (+3.8%, +7.6% and +3.6% for 10, 20 and 30 subjects). All 130 tracked gaits, of individuals and averages, cost more than their standard-cost optimum.

**Figure 5.**
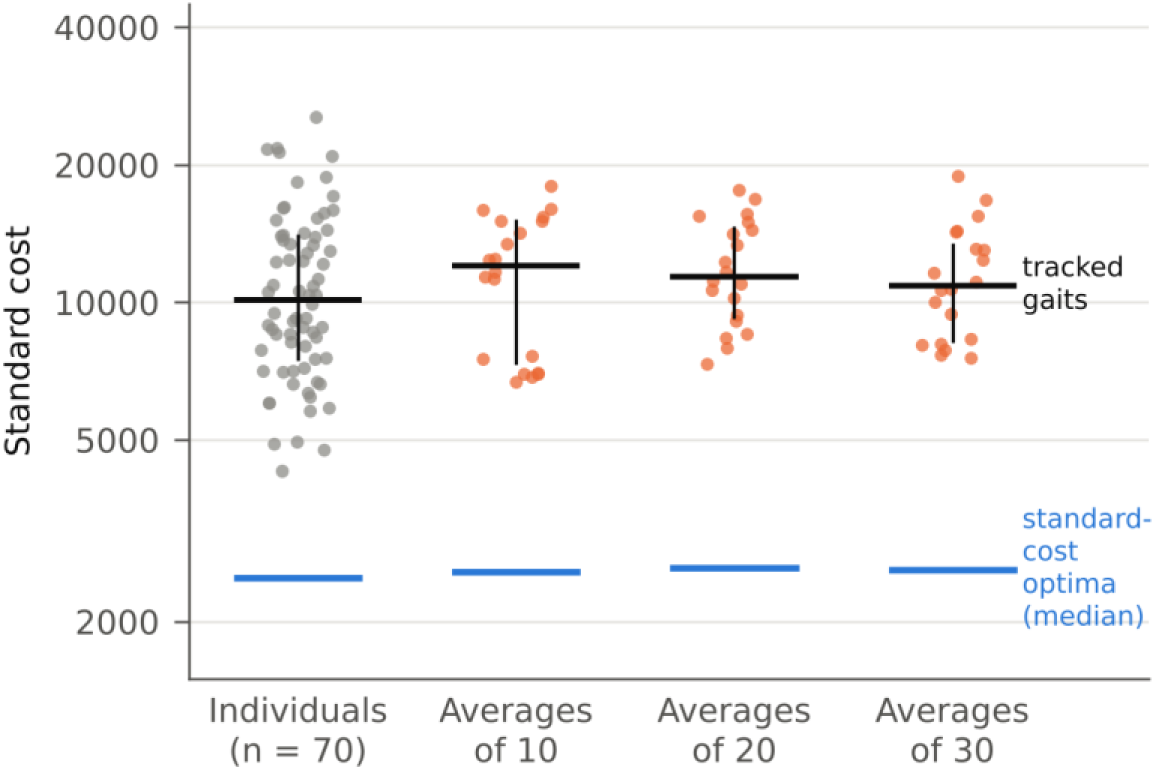
Standard cost of the tracked gaits of individuals and of averages. Individuals and averages of 10, 20 and 30 subjects. Black bars show the median and interquartile range; blue bars show the median standard cost of the standard-cost optima.

On the other hand, the standard-cost optimum of an average had a better tracking score than the members’ optima (Fig. 6). Its tracking error was 30.3% lower (95% CI 26.4 to 34.2%; p < 0.001), and the reduction was the same for 10, 20 and 30 subjects (30.4%, 30.3% and 30.2%). The optima of the averages still matched the data far worse than their tracked gaits, with median tracking error 8.4 against 1.2, worse in all 60 averages.

**Figure 6.**
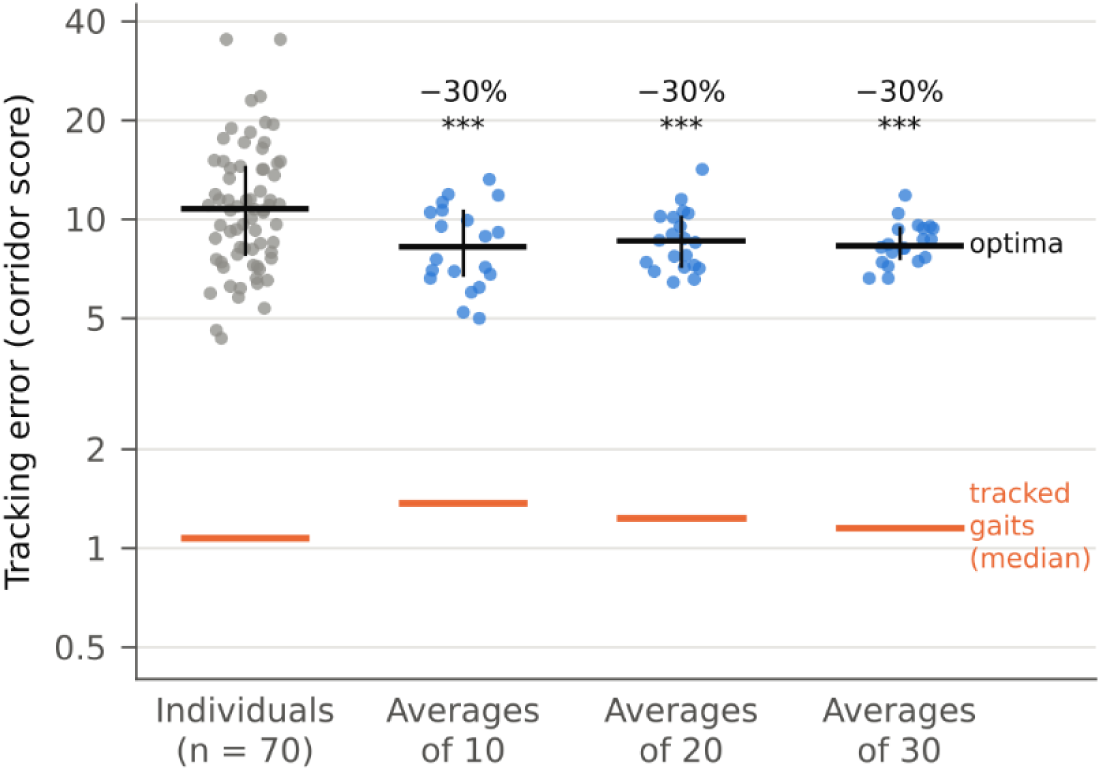
Tracking error of each standard-cost optimum against its own individual or average data. Individuals and averages of 10, 20 and 30 subjects; black bars show the median and interquartile range, orange bars the median tracking error of the tracked gaits. Numbers above each column give the mean difference between the averages’ optima and the mean of their own members’ optima (generalised least squares; *** p < 0.001).

## Discussion

In this work we tested whether optimising the cost functions normally used in predictive gait simulations tracks an individual’s data as closely as the tracked gait obtained with the same model and problem formulation. In all 20 individuals and in both frameworks, the standard-cost optimum tracked the data worse than the tracked gait, and the tracked gait cost more than the optimum. Tuning the weights lowered the tracking error of the optimum by a median 19% in SCONE and 17% in PredSim, but the best of 35 weightings still tracked worse than the tracked gait in 18 of 20 subjects in SCONE and in all 20 in PredSim. The tracked gait here is not a competing predictor, because it was fitted to the measured data. We use it to show that a gait matching the data is dynamically feasible with the same model and constraints. The distance between the optima and the data therefore cannot be explained only by the model’s inability to walk as the person does.

The comparison of joint angles and muscle activity (Tables 2 and 3, Fig. 3 and 4) shows which variables differed most between the optima and the measured data. In PredSim, the joint-angle patterns of the optima were close to the measured ones, but their muscle activity correlated poorly with the EMG. A gait can therefore look plausible in its kinematics while being produced by a different muscle coordination, and comparisons based on kinematics alone may overstate how well a cost describes an individual. In SCONE, the tracked gait followed the EMG poorly, which suggests that the reflex controller limits how closely muscle activity can be tracked and possibly explains the smaller gap between tracking and optimisation in this framework.

We then tested whether the standard cost describes a population better than an individual, with two hypotheses: (1) the tracked gait of an average, built from averaged data and an averaged body, has a lower standard cost than the tracked gaits of its members, and (2) the standard-cost optimum of an average tracks its data better than the members’ optima track theirs. The tracked gaits of averages were no cheaper than those of their members, so hypothesis (1) was not supported. The standard-cost optimum of an average body/data had a 30% lower tracking error than its members’ standard-cost optima had with respect to their own data, confirming hypothesis (2).

This latter result may explain why standard costs have been considered adequate in the literature on predictive gait simulation, where costs ranging from metabolic energy [1] and fatigue-like criteria [2] to weighted combinations of energy, effort, smoothness and stability [31, 32] were evaluated against group data with generic models, which represent an average person [33].

Our analysis shows that standard-cost optimisation replicates average data better. A likely reason is that gait patterns are unique to each person [17], and averaging removes the individual features that the cost cannot produce while keeping the shared ones. In this view, part of the agreement between predictions and data reported in earlier studies may come from averaging, and not from a cost that describes how people control gait. Averaging did not lower the standard cost of the tracked gait, however, and even for averages the standard-cost optimum tracked the data far worse than the tracked gait (median tracking error 8.4 against 1.2). This shows that population data are more similar to the gait that can be produced by optimising a standard cost, but the averaged gait is still not an optimum of the standard cost.

Since a generic cost describes average data better than individual data, a subject-specific cost is likely necessary if we want to predict an individual’s gait. Inverse optimal control (IOC) has been proposed as a way to identify such a cost by estimating the weights of a cost function formulation from measured movement, typically through a bilevel formulation where an inner optimal control problem predicts the gait for a given set of weights, and an outer optimisation adjusts the weights until the predicted gait best matches the measured data [14, 24]. In IOC formulations, identified weights differed between subjects [15] and with walking speed [16], and the fitted gaits were shown to match the data except at the ankle [24].

The match between data and prediction in IOC is still looser than tracking. In previous work [24] the joint-angle error of the IOC fit was 0.90 SD, against 0.36 SD for tracking with the same model. With a planar reflex model similar to the one we used here, a cost tuned to group-mean data gave an RMSE of 2.15 SD, against 1.15 SD for tracking [5]. These differences in joint angles are close in magnitude to the ones we observed between the best weighting and the tracked gait in both PredSim and SCONE, while none of the cited studies tracked the EMG data.

Later IOC methods added task-level terms such as trunk posture and centre-of-mass motion [34], replaced the weighted sum with a Chebyshev scalarisation [35] or allowed the weights to vary over time, in jumping [36]. All of these methods assume that the measured movement is an optimum of the cost for some weighting of the chosen terms. Our weight grid is a coarse IOC over two weights, with the tracking error as the identification criterion. As in Clever et al. [15], the best weighting differed between subjects and also differed between frameworks and was never, for any of the subjects in either framework, the default weighting.

A finer or wider weight search would probably have lowered the tracking error further, but even the optimal weighting of these terms would probably not reach the tracking error of the tracked gait. First because the optima of all weightings were cheap in cost in both formulations. In fact the best weightings cost only 6% (SCONE) and 16% (PredSim) more than the standard-cost optimum whereas the tracked gaits cost 3.5 times as much. Reweighting the cost function terms therefore shifted the optimum among lower cost gaits, but none of these tracked the data as closely as the tracked gait. Second, IOC can only return gaits that are optimal for some weighting. Minimising a weighted sum with positive weights always returns a Pareto-optimal gait, that is, a gait that no other gait improves on every cost term at once [37]. In PredSim, the standard-cost optimum was lower than the tracked gait on every cost term in 16 of 20 subjects, so no positive weighting of these terms could make the tracked gait optimal for those subjects. Tracking has no such restriction, because it searches all dynamically feasible gaits, which may be gaits that are dominated and not optimal.

This raises the question of whether optimisation, which returns a Pareto-optimal solution, is the right generative principle in predictive modelling of gait. Optimality principles explain many features of human motor control. People walk at speeds that minimise energy per unit distance [38], make movements that minimise endpoint variance [39], and weigh effort when choosing how to move [40]. Optimal feedback control also explains why variability is larger in directions that do not affect the task [41, 42]. Predictive simulations use optimality more narrowly, as the minimum of a cost chosen in advance, whether it is solved by collocation, reflex control or reinforcement learning [43].

Optimality has several limits as a basis for predictive simulation. First, errors in the model can raise the cost of the tracked gait, because a model that differs from the person needs extra muscle effort to follow the person’s data. Model and cost errors are also hard to discern. In earlier predictive simulations, the realism of stance knee flexion depended on the cost [2], on the initial kinematics imposed [1] and on whether the toes were modelled [44]. It is therefore hard to separate the limits of the cost from those of the model [33].

Second, even if an individual’s nervous system minimises some cost, that cost cannot be measured directly. Any formulation may therefore include terms that only correlate with the true ones, or may lack terms altogether. Several studies point to terms that standard costs usually omit. Under motor noise, stochastic optimal control predicts co-contraction, which a deterministic effort cost penalises [45]. People also trade energy economy against stability [21, 46] and against overburdening individual muscles [47]. The weights of the cost may also change within a movement or with walking speed [16, 36].

Finally, people may also not minimise a fixed cost at all. Energy cost falls as people adapt to a new walking condition [48], and they re-optimise step frequency within minutes [18], but they do not always find the cheapest gait when free to explore [19, 20], and muscle coordination may be habitual and only good enough [23, 49].

These limits are particularly important in applications where the gait of a specific person needs to be predicted, for example to plan surgery or design an assistive device, or when modelling people with neurological conditions, whose gait may be further from the optimum of a standard cost. After stroke, people walk asymmetrically [50], and walking more symmetrically does not always lower their energy cost [51]. With cognitive decline, gait becomes slower and more variable [52, 53]. For such individuals, subject-specific predictions have been obtained by calibrating the model on the person’s data and keeping the predicted controls close to those of a tracking solution [54]. Another option is to replace hand-tuned costs with priors learned from realistic movement [55].

## Conclusions

To conclude, our work shows that predictive simulations based on the minimisation of a standard cost function generate plausible gait, but they do not describe how a specific person walks. Our results suggest that the limit in representing an individual is not only the model or the representation of control that we use, but also the principle that we use to generate the gait. Tuning the weights of the cost moves the predicted gait closer to an individual’s data, but the solution remains confined to the optima of the chosen cost terms, and far from the data. The apparent adequacy of standard costs in the literature may partly reflect comparisons with group data, which are closer to these optima than individual data are.

The limit of optimality as a generative principle in predictive simulations is particularly important in applications where predictive simulations are expected to inform individual decisions, such as planning surgery, prescribing orthoses or designing assistive devices for people with impaired gait. Predictive simulation of individual gait therefore needs to account for non-optimality. The field should develop formulations in which gait can lie away from the optimum, for example because of habit, incomplete adaptation or priorities other than economy. It should also test these formulations on individuals, not on group averages.

## List of abbreviations

CI: confidence interval
CMA-ES: covariance matrix adaptation evolution strategy
EMG: electromyography
IOC: inverse optimal control
NRMSE: normalised root-mean-square error
SD: standard deviation

## Declarations

### Ethics approval and consent to participate

This study is a secondary analysis of an openly available, anonymised dataset [25]. Ethics approval and informed consent for the original data collection are reported in [25]. No new data were collected from human participants.

### Consent for publication

Not applicable.

### Availability of data and materials

The gait data analysed during the current study are openly available in the dataset of Van Criekinge et al. [25]. The code and simulation results is available upon reasonable request to the corresponding author.

### Competing interests

The authors declare that they have no competing interests.

## Funding

This project was supported by Research Ireland (Grant No. 21/ FFP-P/10228 to GS).

## Authors’ contributions

GS conceived the study. MMA, DM and GS ran the simulations and analysed the data. GS wrote the manuscript. MMA, DM and GS revised the manuscript. All authors approved the manuscript.

## Notes

### Competing Interest Statement

The authors have declared no competing interest.

